# EnhancerBD identifing sequence feature

**DOI:** 10.1101/2024.03.05.583459

**Authors:** Yi Wang

## Abstract

Deciphering the non-coding language of DNA is one of the fundamental questions in genomic research. Previous bioinformatics methods often struggled to capture this complexity, especially in cases of limited data availability. Enhancers are short DNA segments that play a crucial role in biological processes, such as enhancing the transcription of target genes. Due to their ability to be located at any position within the genome sequence, accurately identifying enhancers can be challenging. We presented a deep learning method (enhancerBD) for enhancer recognition. We extensively compared the enhancerBD with previous 18 state-of-the-art methods by independent test. Enhancer-BD achieved competitive performances. All detection results on the validation set have achieved remarkable scores for each metric. It is a solid state-of-the-art enhancer recognition software. In this paper, I extended the BERT combined DenseNet121 models by sequentially adding the layers GlobalAveragePooling2D, Dropout, and a ReLU activation function. This modification aims to enhance the convergence of the model’s loss function and improve its ability to predict sequence features. The improved model is not only applicable for enhancer identification but also for distinguishing enhancer strength. Moreover, it holds the potential for recognizing sequence features such as lncRNA, microRNA, insultor, and silencer.

## 1. Introduction

Enhancers are short DNA segments that play a crucial role in biological processes, such as enhancing the transcription of target genes. Due to their ability to be located anywhere in the genome sequence, accurately identifying enhancers can be challenging. I have proposed a deep learning-based algorithm for enhancer identification, which I refer to as enhancerBD.

According to recent research, biological sequences have been widely applied in bioinformatics studies, such as DNA, RNA, and protein sequences [1]. Understanding the structural features of the desired sequence from sequence databases is one of the main challenges in biological sequence analysis.Therefore, an effective framework is needed to automatically represent sequence information. In recent years, advanced deep learning methods applied to bioinformatics analysis have shown promising results and advantages in handling sequence data, such as protein function prediction [2], electron transfer protein recognition [3], and variable-length antiviral peptide identification [4]. There is increasing evidence of successful applications in protein prediction and genomic analysis. Additionally, biological sequences like DNA and protein sequences can be considered as textual information, sharing some similarities with human language. As a result, some research groups have applied advanced natural language processing (NLP) techniques to learn useful features from text-formatted biological data and have achieved promising results [5]. Integrating embedding techniques into deep learning enables us to address the representation and extraction of features from biological sequences.

In recent years, language representation models have gained considerable attention in the field of natural language processing due to their remarkable achievements. Among them, Bidirectional Encoder Representations from Transformers (BERT[6]) has been proven to be a simple yet powerful language model, achieving state-of-the-art performance.

BERT (Bidirectional Encoder Representations from Transformers) is a deep learning model. Google introduced it in 2018. BERT is a transformer-based model that is pre-trained on a large corpus of unlabeled text data in a self-supervised manner. It adopts a bidirectional approach, meaning it considers the context from both the left and right sides of a word. This bidirectional training allows BERT to capture the rich semantic relationships between words and greatly improves its language understanding capabilities.

During pre-training, BERT learns to predict missing words within a sentence, a process known as masked language modeling. It also learns to predict whether two sentences are consecutive or randomly selected, a task called next sentence prediction. By training on these tasks, BERT learns to encode contextual information within each word representation, capturing the meaning and relationships between words in a given sentence.

After pre-training, BERT can be fine-tuned on specific downstream NLP tasks such as text classification, named entity recognition, question answering, and machine translation. This allows BERT to adapt its learned representations to the specific task at hand and achieve high performance with less labeled data.

One of the key advantages of BERT is its ability to handle various NLP tasks with a single unified model. It captures both syntactic and semantic relationships within text, making it particularly effective in understanding and generating human-like language. In summary, BERT is a state-of-the-art deep learning model for NLP tasks. It leverages transformer-based architecture, bidirectional training, and pre-training followed by fine-tuning on specific tasks. BERT has greatly advanced the field of NLP by achieving remarkable performance on a wide range of natural language understanding tasks.

DenseNet-121[7] is a deep learning algorithm that belongs to the DenseNet family of models. Huang et al. Introduced it in 2016 and has become popular for various computer vision tasks, such as image classification and object detection. DenseNet-121 is specifically designed to address the vanishing gradient problem in deep neural networks and improve information flow throughout the network.

In DenseNet, each layer is directly connected to all subsequent layers, forming a dense block. This dense connectivity allows for direct information exchange between layers, enabling gradients to flow more easily.

DenseNet-121 consists of multiple densely connected blocks, which are composed of convolutional layers, batch normalization, and ReLU activation functions. Each dense block consists of multiple convolutional layers, where the output of each layer is connected to the outputs of all preceding layers within the same block. This dense connectivity ensures that the subsequent layers have access to all previously learned features, leading to an efficient reuse of feature maps and reducing the risk of information loss.

In addition to dense blocks, DenseNet-121 also includes transition blocks. They consist of a batch normalization layer, followed by a 1x1 convolutional layer and average pooling, which downsamples the feature maps.

DenseNet-121 is composed of four dense blocks and three transition blocks. The input to the network is fed through a convolutional layer, followed by a dense block, a transition block, another dense block, another transition block, another dense block, and a final transition block.

DenseNet-121 has several advantages, including its strong feature reuse capabilities, efficient use of parameters, and computational efficiency. It has achieved excellent performance on various benchmark datasets and is widely used as a backbone architecture in many computer vision applications.

In summary, DenseNet-121 is a deep learning algorithm that addresses the vanishing gradient problem by introducing dense connectivity between layers. It consists of densely connected blocks and transition blocks, which facilitate efficient feature reuse and reduce information loss. DenseNet-121 has proven to be highly effective for image classification tasks and has become a popular choice in the computer vision community.

As critical components of DNA, enhancers can efficiently and specifically manipulate the spatiotemporal regulation of gene transcription. Malfunction or dysregulation of enhancers is implicated in a slew of human pathology[8].

Enhancing the transcription of target genes is one of the major function of enhancer [9,10]. Unlike promoters, enhancers do not need to be located near the transcription start site. [10-12]. Increasing evidence suggests that enhancers play a critical role in gene regulation [13,14]. Enhancers control the expression of genes involved in cell differentiation [15, 16]. They coordinate crucial cellular events such as differentiation [17, 18], maintenance of cell identity [19, 20], and response to stimuli [21-23] by binding with transcription factors [24]. Enhancers are closely associated with inflammation and cancer [25].Therefore, this experiment is designed to use deep learning model software based on natural language processing models to identify enhancer sequences to provide powerful help for biological genetic information analysis.

## 2. DATA

We use iEnhancer-2L study’s dataset to address the DNA enhancer classification problem[26]. In this dataset, the original authors collected enhancer sequences from 9 different cell lines and fragmented them into 200 bp segments. Using CD-HIT to remove DNA sequences with high similarity (>20%) [27]. The final data consists of 1484 enhancers and 1484 non-enhancers in the training set, and 200 enhancers and 200 non-enhancers in the independent set. 742 strong and 742 weak enhancers in the training set and 100 strong enhancers and 100 weak enhancers in the independent set[28].

## 3. Methods

In this paper, I extended the BERT combined DenseNet121 models by sequentially adding the layers GlobalAveragePooling2D, Dropout, and a ReLU activation function. This modification aims to enhance the convergence of the model’s loss function and improve its ability to predict sequence features. The improved model is not only applicable for enhancer identification but also for distinguishing enhancer strength. Moreover, it holds the potential for recognizing sequence features such as lncRNA, microRNA, insultor, and silencer.

### 3.1. Bert processing

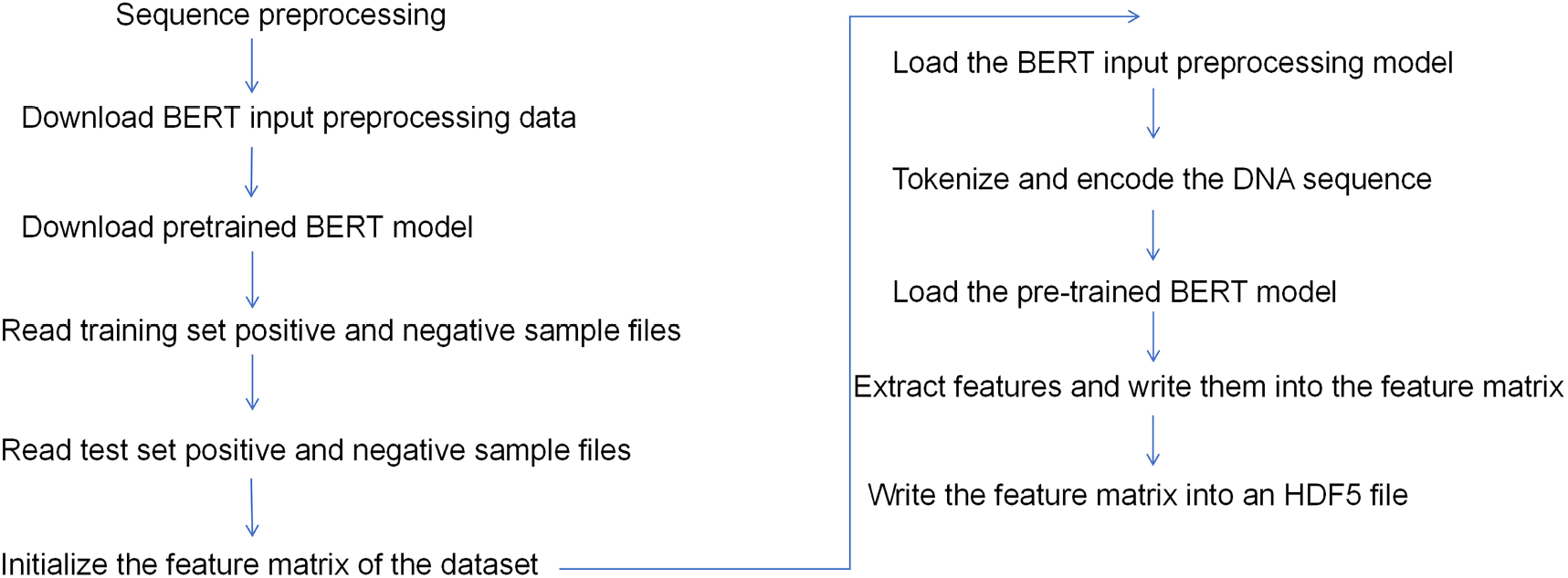

### 3.2. DenseNet121 processing

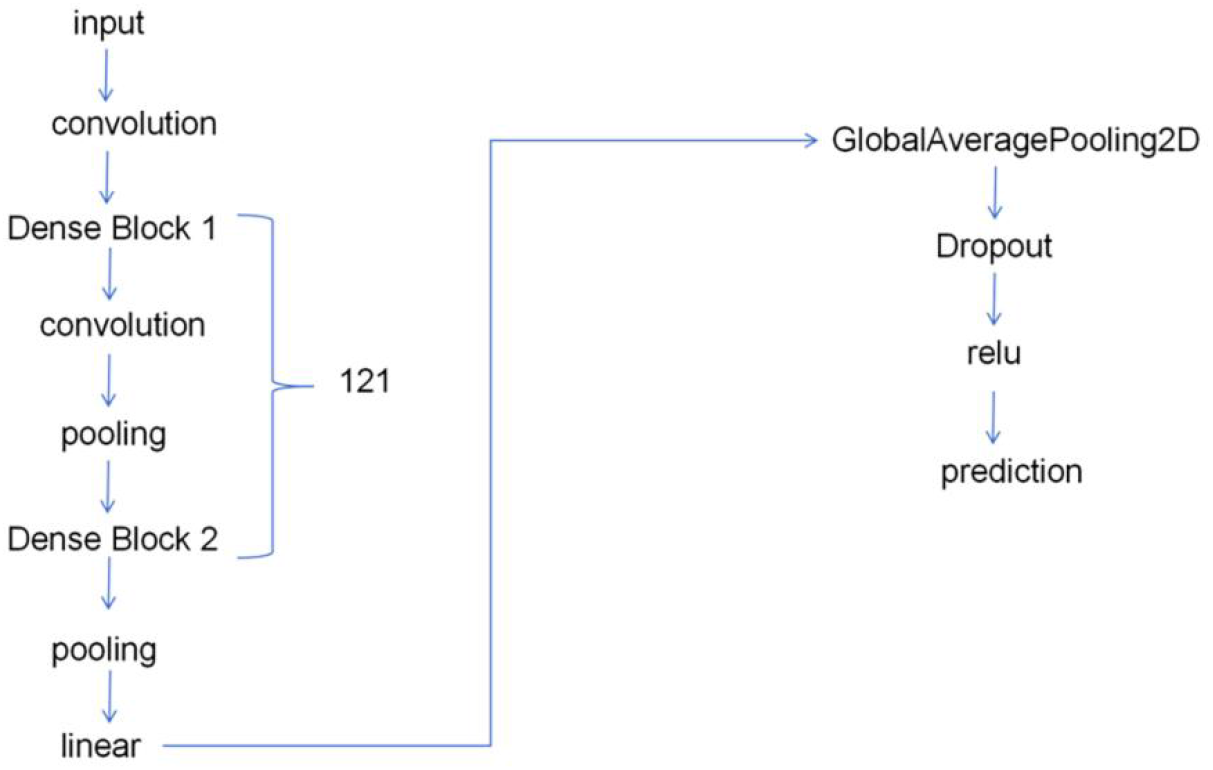

### 3.3. Global Average Pooling 2D

Global Average Pooling 2D (GAP) is a pooling operation commonly used in convolutional neural networks (CNNs) for image analysis tasks. It is applied after the convolutional and activation layers.

GAP operates by taking the average of each feature map along its spatial dimensions (height and width). It helps in capturing the global context and overall summary of the feature maps, making the network more robust to spatial translations and reducing the number of parameters and computations required. This averaging operation helps to maintain more spatial information, making it particularly useful when working with spatially dense features.

After applying GAP, the output feature maps will have a reduced spatial dimension and a depth equal to the number of channels. These features can then be connected to fully connected layers or used as input for subsequent layers in the network for classification or further processing. It helps reduce computational complexity, extract global context, and maintain spatial information in the network.

### 3.4. Dropout Layer

Hinton et al. proposed dropout [29] and is a concept for training deep neural networks. During the training process, a certain proportion of neurons are randomly dropped out, while all neurons are still used during the prediction process [30]. Dropout serves two purposes: accelerating the training of deep neural networks and reducing overfitting. In this experiment, the Dropout parameter is set to 0.1 to distinguish it from higher parameter settings and makes the loss function converge.

### 3.5. Relu

The ReLU function has several advantages. Firstly, it is computationally efficient, as the function is simple and does not require complex calculations. Secondly, it helps alleviate the vanishing gradient problem, which can occur with other activation functions in deep networks. Lastly, ReLU has been empirically shown to be useful in improving the overall performance and convergence speed of deep neural networks in various applications, including image classification, object detection, and natural language processing. In this experiment, a relu activation function layer is added after densenet121 to make the loss function converge.

### 3.6. Performance Measures

The overall prediction accuracy (ACC), sensitivity (SN), specificity (SP), and Matthew’s correlation coefficient (MCC) are used to quantitatively measure the prediction performance. which are defined as

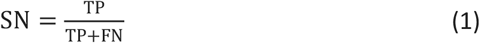

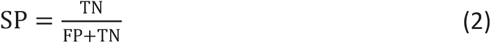

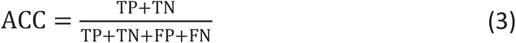

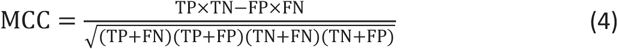

In the equation, TP represents the true positive sample count, FN represents the false negative sample count, FP represents the false positive sample count, and TN represents the true negative sample count. SN, SP, ACC are between 0 and 1. MCC ranges from -1 to 1. The higher the values of SN, SP, ACC, and MCC, the better the performance.

Receiver Operating Characteristic (ROC) curve is a commonly used method to evaluate the performance of binary classification methods. The ROC curve is plotted by graphing the True Positive Rate (TPR) and False Positive Rate (FPR) at different thresholds:

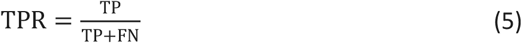

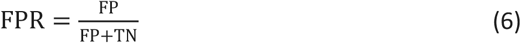

The Area Under the ROC Curve (AUC) ranges from 0 to 1. If AUC equals 1, it indicates a perfect prediction. AUC = 0.5 represents random prediction, while AUC = 0 implies an opposite prediction.

## 4. Result

In this paper, I extended the BERT combined DenseNet121 models by sequentially adding the layers GlobalAveragePooling2D, Dropout, and a ReLU activation function. This modification aims to enhance the convergence of the model’s loss function and improve its ability to predict sequence features. The improved model is not only applicable for enhancer identification but also for distinguishing enhancer strength. Moreover, it holds the potential for recognizing sequence features such as lncRNA, microRNA, insultor, and silencer.

### 4.1. Comparison with State-of-the-Art Methods

This experiment uses the scoring method similar to the study by Guohua Huang et al.[28]. We compared the scoring results of nineteen software including ours and the enhacnerBD performance is the state-of-the-art. All enhancerBD evaluation results are full marks(table1).

**Table 1.**
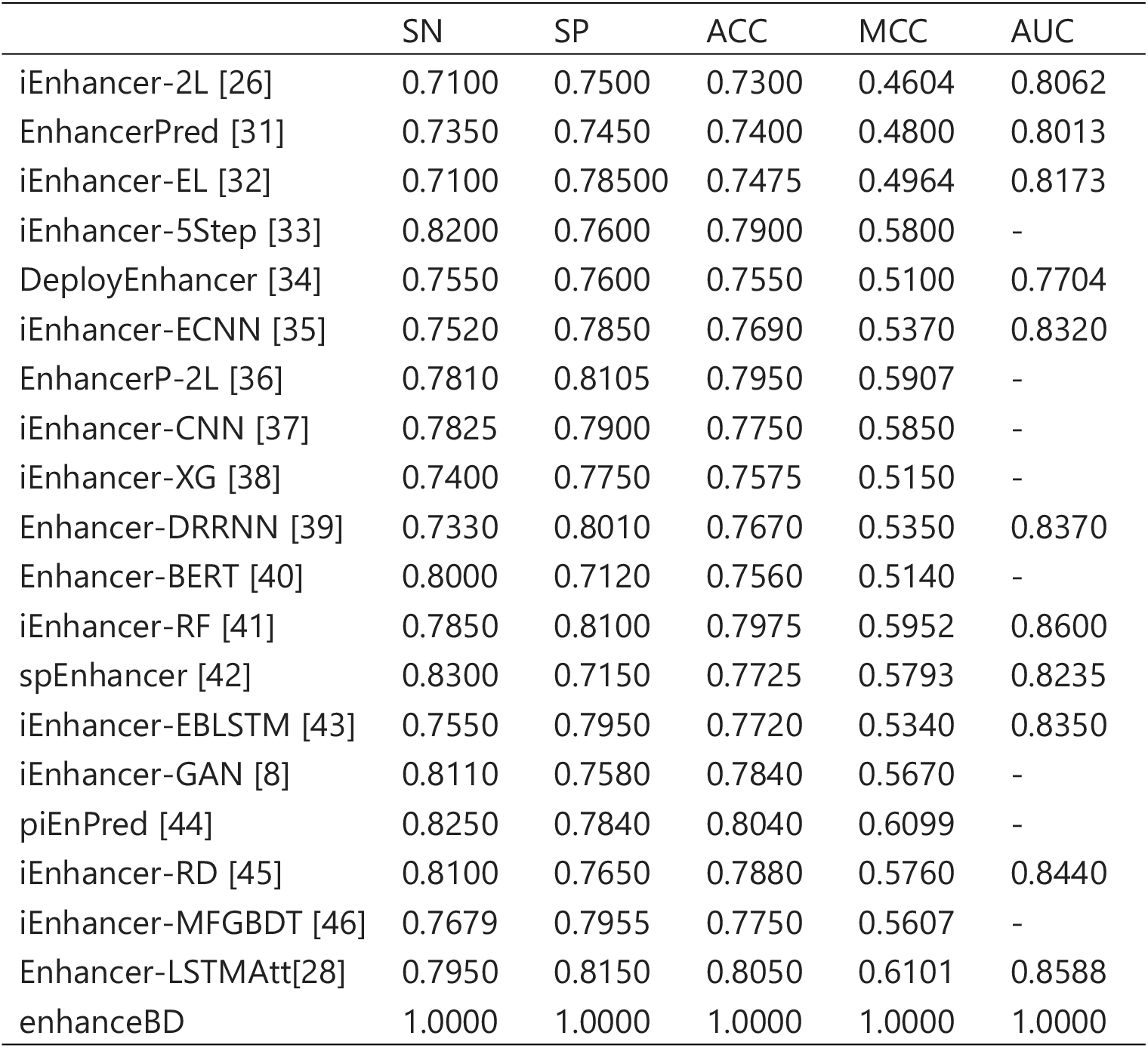
Comparison with state-of-the-art methods by independent test. The first eighteen detection results are sourced from the previous study conducted by Guohua Huang et al. SN: Sensitivity; SP: Specificity; ACC: Accuracy; MCC: Matthew’s Correlation Coefficient; AUC: Area Under the Curve.

|  | SN | SP | ACC | MCC | AUC |
| --- | --- | --- | --- | --- | --- |
| iEnhancer-2L [26] | 0.7100 | 0.7500 | 0.7300 | 0.4604 | 0.8062 |
| EnhancerPred [31] | 0.7350 | 0.7450 | 0.7400 | 0.4800 | 0.8013 |
| iEnhancer-EL [32] | 0.7100 | 0.78500 | 0.7475 | 0.4964 | 0.8173 |
| iEnhancer-5Step [33] | 0.8200 | 0.7600 | 0.7900 | 0.5800 | - |
| DeployEnhancer [34] | 0.7550 | 0.7600 | 0.7550 | 0.5100 | 0.7704 |
| iEnhancer-ECNN [35] | 0.7520 | 0.7850 | 0.7690 | 0.5370 | 0.8320 |
| EnhancerP-2L [36] | 0.7810 | 0.8105 | 0.7950 | 0.5907 | - |
| iEnhancer-CNN [37] | 0.7825 | 0.7900 | 0.7750 | 0.5850 | - |
| iEnhancer-XG [38] | 0.7400 | 0.7750 | 0.7575 | 0.5150 | - |
| Enhancer-DRRNN [39] | 0.7330 | 0.8010 | 0.7670 | 0.5350 | 0.8370 |
| Enhancer-BERT [40] | 0.8000 | 0.7120 | 0.7560 | 0.5140 | - |
| iEnhancer-RF [41] | 0.7850 | 0.8100 | 0.7975 | 0.5952 | 0.8600 |
| spEnhancer [42] | 0.8300 | 0.7150 | 0.7725 | 0.5793 | 0.8235 |
| iEnhancer-EBLSTM [43] | 0.7550 | 0.7950 | 0.7720 | 0.5340 | 0.8350 |
| iEnhancer-GAN [8] | 0.8110 | 0.7580 | 0.7840 | 0.5670 | - |
| piEnPred [44] | 0.8250 | 0.7840 | 0.8040 | 0.6099 | - |
| iEnhancer-RD [45] | 0.8100 | 0.7650 | 0.7880 | 0.5760 | 0.8440 |
| iEnhancer-MFGBDT [46] | 0.7679 | 0.7955 | 0.7750 | 0.5607 | - |
| Enhancer-LSTMAtt[28] | 0.7950 | 0.8150 | 0.8050 | 0.6101 | 0.8588 |
| enhanceBD | 1.0000 | 1.0000 | 1.0000 | 1.0000 | 1.0000 |

### 4.2. Identification of Enhancers

The accuracy prediction value of enancerBD in identifying enhancers is 1 (figure1), the loss score is 0 (figure2), and the AUC score is 1 (figure3).

**Figure 1.**
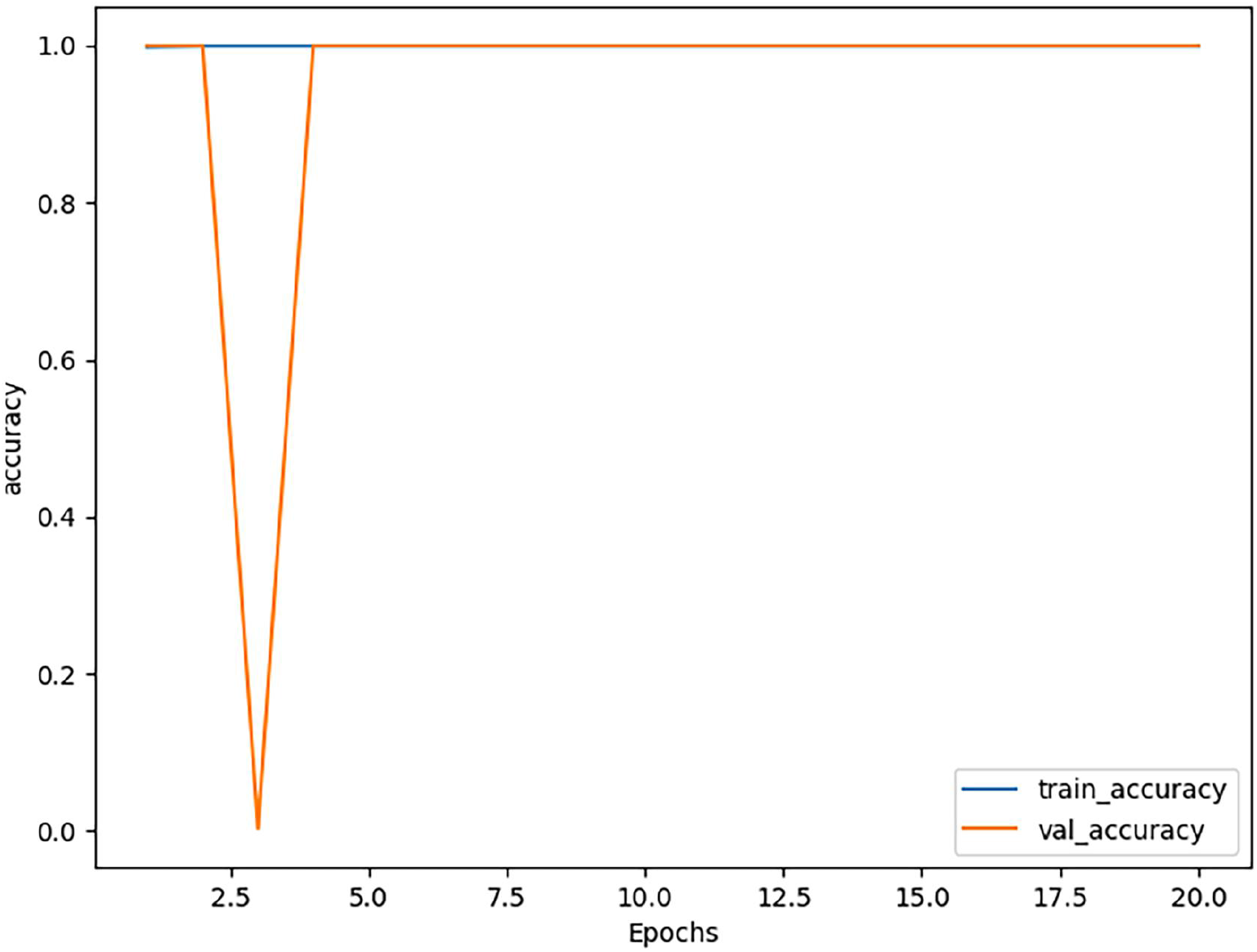
EnancerBD identifies enhancer identity accuracy predictions. “Train” is the training set score and “val” is the test set score. The “epochs” on the abscissa are the generations of training and testing; the “accuracy” on the ordinate is the accuracy score.

**Figure 2.**
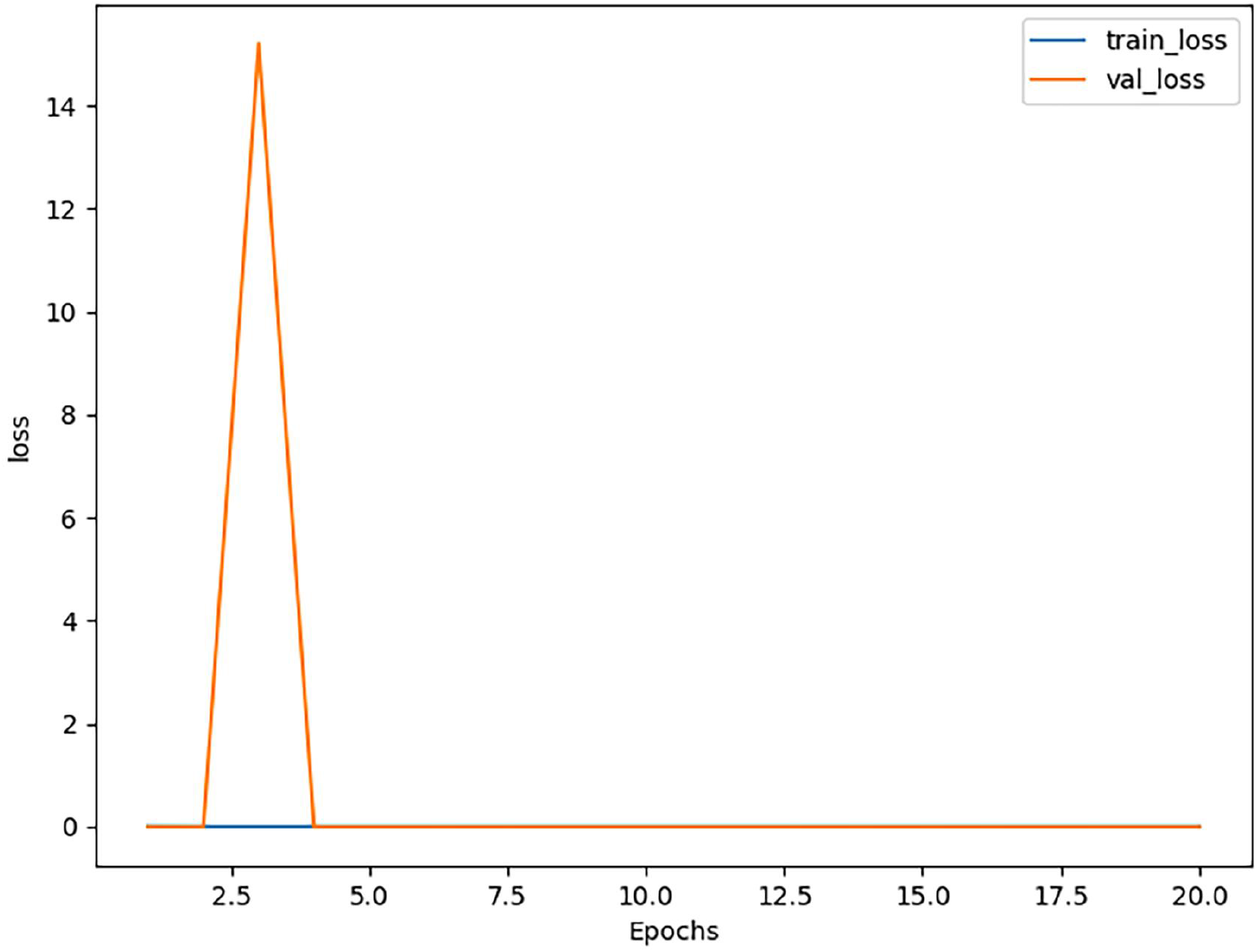
EnancerBD Loss function for identifying enhancers. “Train” is the training set score and “val” is the test set score. The “epochs” on the abscissa are the generations of training and testing; the “loss” on the ordinate is the score of the loss function.

**Figure 3.**
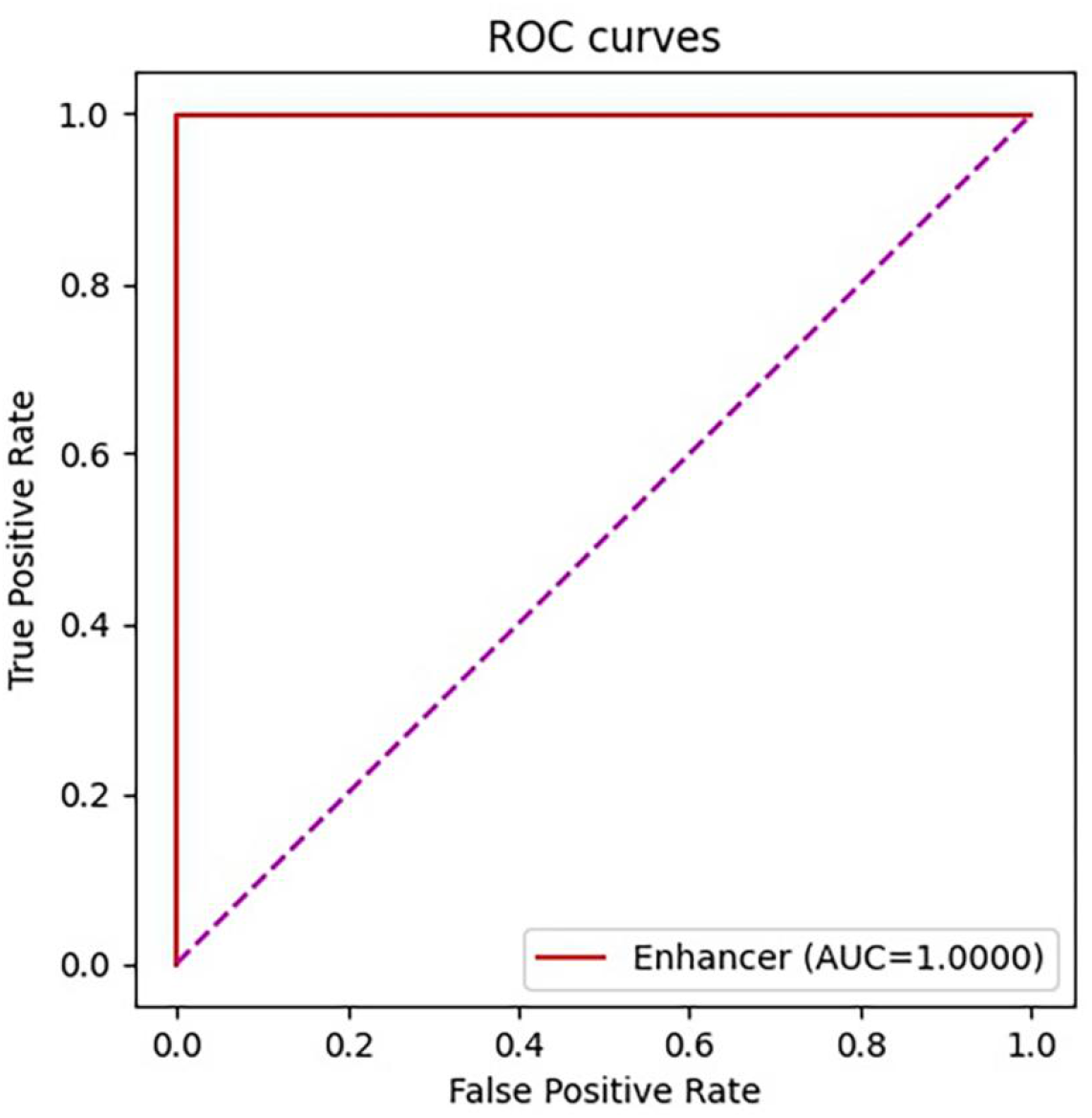
The AUC curve of EnancerBD identifies enhancers.The purple line is the baseline of ROC curve, named random guess line.

### 4.3. Distinguish the strength of enhancers

The prediction accuracy of enancerBD for enhancer strength is 1 (figure4), the loss score is 0 (figure5), and the AUC score is 1 (figure6). Not large enough training dataset maybe the cause of the slight difference between the scores of the training set and test set.

**Figure 4.**
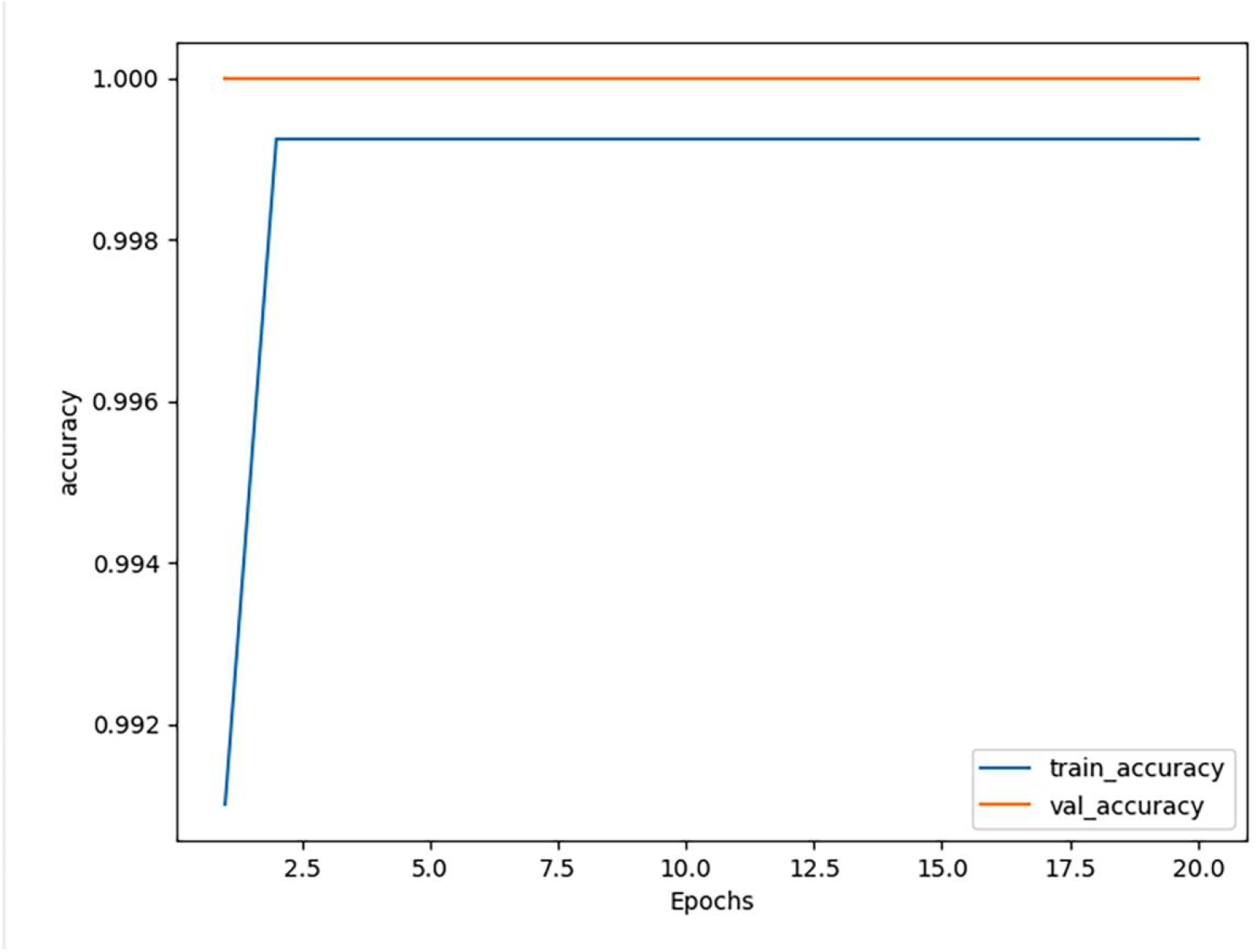
The prediction accuracy of enhancer strength by enancerBD. “Train” is the training set score and “val” is the test set score. The “epochs” on the abscissa are the generations of training and testing; the “accuracy” on the ordinate is the accuracy score.

**Figure 5.**
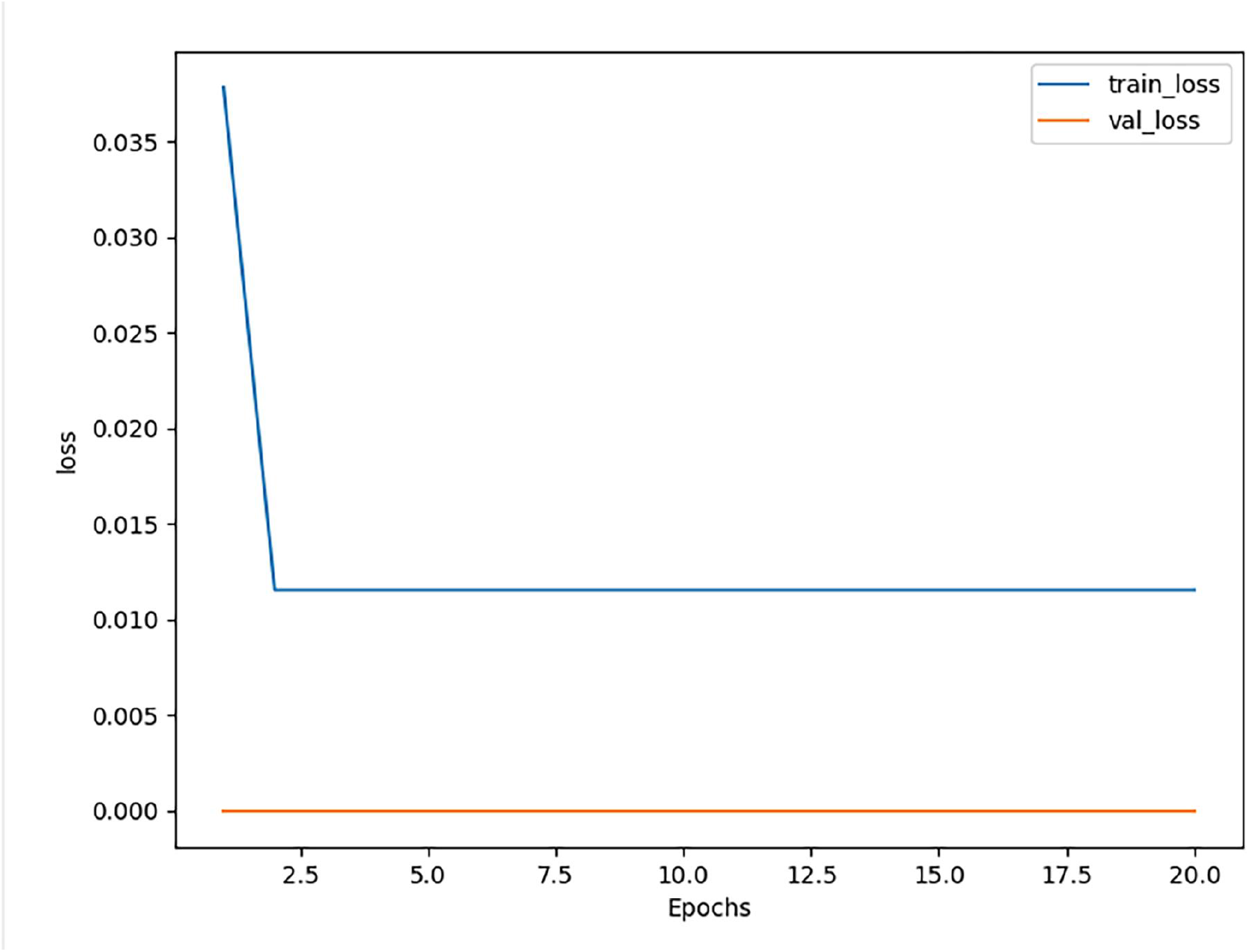
The loss function for predicting the strength of enhancers in enhancerBD. “Train” is the training set score and “val” is the test set score. The “epochs” on the abscissa are the generations of training and testing; the “loss” on the ordinate is the score of the loss function.

**Figure 6.**
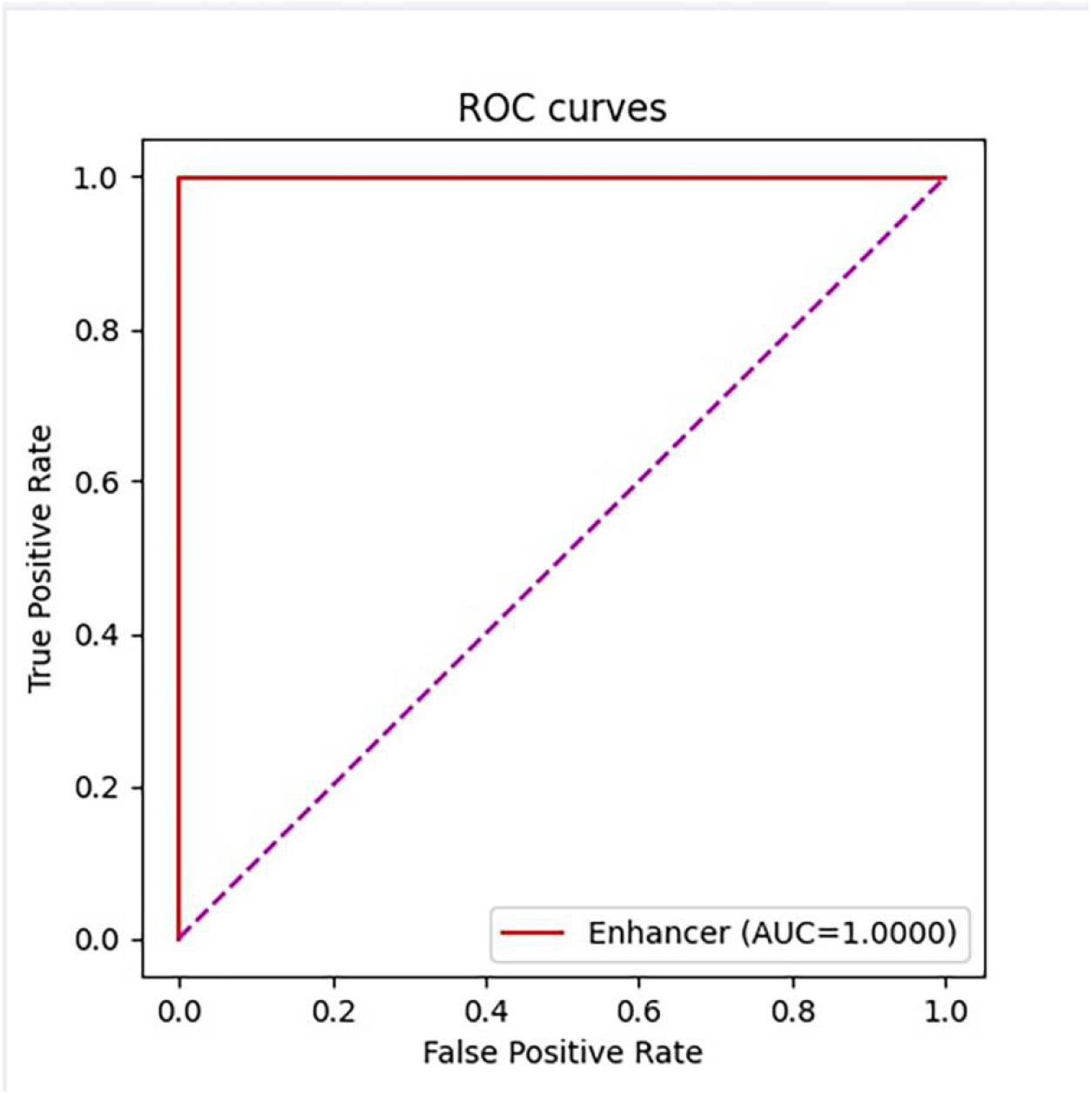
AUC score of enhancer strength prediction by enancerBD.The purple line is the baseline of ROC curve, named random guess line.

### 4.4. Discussion

The software enhancerBD developed in this experiment is the state-of-the-art in the field of identifying enhancers compared with many previous software, and the scores are all full marks. In terms of identifying the strength and weakness of enhancers, it still achieved remarkable results.

The model enhancerBD used is not only applicable for enhancer identification but also for distinguishing enhancer strength. Moreover, it holds the potential for recognizing sequence features such as lncRNA, microRNA, insultor, and silencer.

## 5. Conclusions

As crucial regulatory elements of DNA, enhancers play a vital role in gene transcription and are associated with a range of diseases. Accurate identification of enhancers and their strength can contribute to uncovering potential mechanisms underlying enhancer-related biological processes and disease progression.The enhancer identification software, enhancerBD, designed in this study has achieved state-of-the-art performance in previous evaluations. Whether it is distinguishing enhancers from non-enhancers or assessing the strength of enhancers, the software consistently shows detection results with an AUC of up to 1. Such high-performance detection capabilities enable the software to be widely applied in disease research, providing significant assistance in understanding gene sequences. In this paper, I extended BERT combined DenseNet121 models by sequentially adding the layers GlobalAveragePooling2D, Dropout, and a ReLU activation function. This modification aims to enhance the convergence of the model’s loss function and improve its ability to predict sequence features. The improved model is not only applicable for enhancer identification but also for distinguishing enhancer strength. Moreover, it holds the potential for recognizing sequence features such as lncRNA, microRNA, insultor, and silencer.

## Author Contributions

Conceptualization, Y.W.; methodology, Y.W.; software, Y.W.; validation, Y.W.; formal analysis, Y.W.; investigation, Y.W.; resources, Y.W.; data curation, Y.W.; writing—original draft preparation, Y.W.; writing—review and editing, Y.W.; project administration, Y.W.; All authors have read and agreed to the published version of the manuscript.

## Institutional Review Board Statement

Not applicable.

## Informed Consent Statement

Not applicable

## Data Availability Statement

(We will make the code public on github when the article is published)

